# Leveraging Descriptor Learning and Functional Map-based Shape Matching for Automatic Landmark Acquisition

**DOI:** 10.1101/2024.05.22.595350

**Authors:** Oshane O. Thomas, A. Murat Maga

## Abstract

Geometric morphometrics is widely employed across the biological sciences for the quantification of morphological traits. However, the scalability of these methods to large datasets is hampered by the requisite placement of landmarks, which can be laborious and time consuming if done manually. Additionally, the selected landmarks embody a particular hypothesis regarding the critical geometry pertinent to the biological inquiry at hand. Modifying this hypothesis lacks flexibility, necessitating the acquisition of an entirely new set of landmarks on the entire dataset to reflect any theoretical adjustments. In our research, we investigate the precision and accuracy of landmarks derived from the comprehensive set of functional correspondences acquired through the functional map framework of geometry processing. We use a deep functional map network to learn shape descriptors that effectively yield functional map-based and point-to-point correspondences between the specimens in our dataset. We then interrogate these maps to identify corresponding landmarks given manually placed landmarks from the entire dataset. We assess our method by automating the landmarking process on a dataset comprising mandibles from various rodent species, comparing its efficacy against MALPACA, a cutting-edge technique for automatic landmark placement. Compared to MALPACA, our model is notably faster and maintains competitive accuracy. The Root Mean Square Error (RMSE) analysis reveals that while MALPACA generally exhibits the lowest RMSE, our models perform comparably, especially with smaller training datasets, suggesting strong generalizability. Visual evaluations confirm the precision of our landmark placements, with deviations remaining within an acceptable range. These findings underscore the potential of unsupervised learning models in anatomical landmark placement, providing a viable and efficient alternative to traditional methods.

## 1. Introduction

Landmark-based morphometrics constitutes a critical methodology in biological research, enabling precise quantitative analysis of shape and size variation in organisms (Adams et al., 2004; Webster and Sheets, 2010; Mitteroecker and Schaefer, 2022). Central to studies in developmental, evolutionary, and functional morphology, landmark-based morphometrics allows researchers to rigorously test hypotheses concerning genetic and environmental impacts on morphological traits (Dunn and Avery, 2021; Hobbs et al., 2021). Such studies deepen our understanding of phenotypic plasticity and adaptation, and are indispensable in phylogenetic analyses, helping trace morphological changes across evolutionary timelines and clarifying taxonomic relationships (Lawing and Polly, 2010).

As morphological data increasingly become pivotal in integrative biology, automating measurement and standardizing landmark placement is becoming essential (Porto et al., 2021; Thomas et al., 2023). This automation is crucial for supporting large-scale studies and meta-analyses that require consistent data quality from diverse sources. Emerging techniques, particularly those leveraging advancements in machine learning and unsupervised algorithms, offer substantial improvements in processing efficiency and robustness (MacLeod et al., 2010; Macleod, 2017). Such technologies enable researchers to manage larger datasets with minimal bias and enhanced throughput, significantly advancing the field (Adams and Otárola-Castillo, 2013; Boyer et al., 2014; Maga et al., 2015).

The automation of anatomical landmark collection not only streamlines the investigative process but also enhances precision and reproducibility, which are vital for advancing our understanding of complex biological structures. Automated systems reduce the subjectivity and inconsistencies typical of manual landmark identification, ensuring that findings are robust and universally comparable. This integration saves valuable resources and augments the capabilities of computational biology, enabling intricate exploration of patterns and relationships within and across species. Furthermore, these advancements improve the accessibility of detailed morphological analyses, promoting a more inclusive and comprehensive approach to anatomical research (Richtsmeier et al., 2002, 2005).

However, automatic landmarking of biological specimens introduces significant challenges that impact the reliability and validity of morphometric analyses. Biological specimens often exhibit considerable variability due to genetic, developmental, and environmental factors, challenging algorithms to consistently identify landmarks across diverse samples (Devine et al., 2019). These issues are compounded by the potential misinterpretation of subtle anatomical features by automated systems, which could skew morphological analyses. The development of robust automatic landmarking systems thus hinges on the availability of extensive, precisely annotated datasets essential for training machine learning models to accurately generalize across the natural diversity found within and between species. Addressing these challenges is critical to enhancing the precision and utility of automatic landmarking tools in anatomical and evolutionary biology research (Slice, 2006, 2007; Adams and Otárola-Castillo, 2013).

The objective of this study is to develop and evaluate a deep learning-based algorithm specifically designed for the automatic landmarking of morphological specimens represented as 3D polygon meshes. Our methodology leverages advanced geometry processing techniques to establish correspondences between biological shapes using the Functional Map (FMap) framework. Geometry processing is a field focused on the acquisition, representation, processing, analysis, and visualization of geometric data. The FMap framework is an approach within this field that represents maps between shapes as transformations of functions rather than direct point-to-point correspondences. This method involves using the eigenfunctions of the Laplace-Beltrami operator to create a compact and stable map representation, which simplifies the application of constraints and the inference process. The framework is particularly useful for tasks such as shape matching, segmentation transfer, and the joint analysis of shape collections, allowing for efficient and accurate manipulation of complex geometries. Applications of this framework include improving shape matching methods, transferring annotations between shapes, and facilitating the study of anatomical structures through detailed comparative analysis (Ovsjanikov et al., 2012, 2017; Ovsjanikov, 2016; Thomas et al., 2023).

Our research builds on the foundational work of the MorphVQ pipeline introduced by (Thomas et al., 2023). MorphVQ takes advantage of the FMap framework for biological shape analysis, and was intended for landmark-free representation of shape difference as it uses shape operators (latent shape variables) to represent differences in morphology within a collection of bone shapes. Unlike the original MorphVQ, our approach aims to generate accurate and precise dense point-to-point correspondences from FMap correspondences, facilitating the acquisition of landmarks. Thomas et al. 2023 highlighted the efficacy of an FMap-based pipeline for biological shape analysis, showing how this approach can successfully characterize both intra- and inter-group variation for diverse research applications. Notably, the efficacy of MorphVQ was evaluated by comparing the error between landmarks placed by the algorithm and those ground-truth landmarks collected by an expert, demonstrating the pipeline’s ability to generate precise and accurate landmarks.

This paper advances the field of morphometric analysis by introducing significant enhancements to the MorphVQ pipeline, improving both the efficiency and accuracy of automatic landmarking in biological specimens. In addition to retooling MorphVQ primarily for landmark acquisition, we enhance the published MorphVQ model (Thomas et al., 2023) in three critical areas: firstly, by eliminating the necessity for the rigid pre-alignment of specimens; secondly, by reducing the time required to train the model; and finally, by improving the quality of the FMaps and the precision of point-to-point correspondences obtained. To achieve these objectives, we have integrated three key algorithmic innovations into our framework. First, our model incorporates the orientation-preserving properties of the newly developed Complex FMap method (Donati et al., 2022). Second, the use of the DiffusionNet model for descriptor learning improves the robustness and adaptability of functional mappings across varying mesh resolutions and surface complexities (Sharp et al., 2020). Third, enforcing spatial and spectral cycle consistency during the Deep Functional Maps (DFMaps) network training process ensures more accurate and bijective point-to-point mappings, reducing the need for post-processing refinement and improving generalization performance across diverse biological datasets (Lähner et al., 2017; Litany et al., 2017; Halimi et al., 2018; Sun et al., 2023). Collectively, these innovations not only augment the capabilities of MorphVQ in morphometrics but also expand its potential for large-scale, high-throughput studies, fostering deeper insights into the evolutionary and developmental dimensions of organismal biology.

The remainder of this paper details the methods and subsequent results that underpin our investigation into the enhanced MorphVQ pipeline. We commence with a comprehensive description of the experimental setup, wherein the MorphVQ model is meticulously evaluated using a mouse mandible dataset—a choice motivated by its relevance and frequent use in morphometric research. This dataset provides a robust basis for assessing the precision and effectiveness of our landmark placement methodology under realistic conditions.

In a joint results and discussion section, we present a thorough analysis of our findings, primarily through an error study that quantifies the landmark placement accuracy of our method. We achieve this by comparing the root mean square error (RMSE) of landmarks estimated by our enhanced MorphVQ model against ground-truth landmarks, with a particular focus on how our model’s performance stands against MALPACA, the current state-of-the-art method in automatic landmark placement. This comparative approach not only highlights the improvements our method offers over existing technologies but also provides a clear, empirical basis to discuss the implications of our results for the broader field of morphological research.

## 2. Materials and Methods

### 2.1 Dataset Description & Polygon Mesh Preprocessing

This study uses a dataset originally characterized by (Maga et al., 2015), comprising 425 3D hemi-mandible models generated from the microCT scans of mouse heads. This dataset was derived from a backcross ([A/J x C57BL/6J] x A/J) of two commonly used inbred laboratory strains. All samples were collected at 28-day postnatally. Briefly, heads of these mice were scanned at 18 micron resolution via microCT. For specifics of the backcross design, husbandry and 3D imaging we refer the reader to Maga et al, 2015, and specifically for generation of the mandibles and associated mandibular landmarks to Navarro and Maga 2016. In this project, data derived from these papers underwent a comprehensive preprocessing pipeline using several modules within the open-source platform, 3D Slicer and its SlicerMorph extension (Fedorov et al., 2012; Rolfe et al., 2021). This initial phase involved the segmentation of the mandibles from the surrounding cranial structures. Following segmentation, we extracted surface polygon models (triangular meshes) from each specimen. These models were then subjected to a cleaning process within 3D Slicer, aimed at refining the data for subsequent analytical steps. We employed the Meshfix python library to address and rectify any deficiencies in the polygon models, such as gaps or non-manifold geometries (Attene, 2010). Meshfix facilitates the seamless filling of holes and ensures that each model adheres to manifold criteria, thereby enhancing the integrity and utility of the extracted models for our morphological analyses.

### 2.2 Ground-Truth Landmarks

The morphometric analysis incorporated in this study uses a set of landmarks delineated and collected in (Navarro and Maga, 2016), comprising thirteen points placed on each specimen’s mandible, annotated on the hemi-mandibles derived from the microCT images (Figure 2). This selection of landmarks aligns with the classical set of landmarks traditionally employed in genetic studies of mouse mandibles (Klingenberg et al., 2001; Workman et al., 2002; Leamy et al., 2008, 2015; Suto and Jun-ichi, 2009; Burgio et al., 2012; Boell, 2013; Boell et al., 2013), offering a robust framework for comparative analysis. To ensure the highest degree of measurement accuracy and reproducibility, each specimen was landmarked twice by a single person. The resultant measurements were then averaged, providing a refined estimate that mitigates potential intraobserver errors inherent in single measurements.

### 2.3 Implementation: Unsupervised Training Scheme and Loss Criterion

Initially, the polygon models representing each specimen were uniformly scaled to unit area. For FMap estimation within our network, we derived the stiffness and mass matrices for each mesh using the cotangent weighting scheme (Ovsjanikov et al., 2016). This critical step facilitated the computation of the Laplace-Beltrami Operator (LBO), executed through Eigen decomposition (Ovsjanikov et al., 2012, 2017; Ovsjanikov, 2016). The process yielded essential eigenvectors and eigenvalues, forming the basis for our analytical endeavors. We also computed the gradient operator for each mesh as complex sparse matrices for use in descriptor learning and complex functional map estimation within our network. These variables, in addition to the vertices of each mesh, were retained for training.

For the unsupervised training of our model, we initially separated the dataset of 425 specimens into two principal groups. The primary group, comprising 322 specimens, was designated for the unsupervised training phase. The secondary group, containing 103 specimens, was specifically reserved for post-training inference to evaluate the model’s ability to generalize to unseen data. These specimens were intentionally excluded from the training phase to provide a stringent test of the model’s generalizability to unseen data.

Further, we subdivided the primary group of 322 samples into two smaller datasets, each consisting of 30 randomly selected specimens drawn without replacement. These subsets, referred to as Dataset 30A and Dataset 30B, were created to investigate the model’s performance under conditions of limited training data, which is common in case of non-model biological systems. This approach allows us to assess the efficacy of unsupervised learning when applied to smaller datasets and to understand the model’s capability in generalizing from minimal training examples to novel data as well.

Our MorphVQ DFMap-based shape matching framework is inherently pairwise in its operation, i.e., every sample is matched to every other sample. Consequently, we structured both the training and inference processes to accommodate this design by ensuring that each pair of shapes within the dataset was analyzed bidirectionally. Specifically, for each of the four datasets, every possible pair was processed from each shape acting as the source to the other as the target, and vice versa. An epoch within our framework was defined as a complete cycle through all such bidirectional pairings. Consequently, the computational complexity of completing one epoch is denoted by *O(n^2^)*, reflecting the intensive pairwise calculations required across the entire dataset.

We developed our model using Pytorch, as outlined by (Li et al., 2020), and incorporated the DFMap component of the MorphVQ framework from Sun et al., 2023. The core of our architecture, the DiffusionNet function approximator, comprised 12 DiffusionNet Block layers, each with an MLP width of 512. Notably, these layers produced an output of 256 descriptor features in the final dimension. Input to the model was provided as Wave Kernel Signatures (WKS), calculated using the first 128 bases of the Laplace-Beltrami Operator (LBO) of each shape, generating 256 descriptor features for each vertex dimension.

Our training regimen utilized a batch size of 4, while inference operated on a batch size of 1. We used a NVIDIA RTX A6000 GPU to conduct all our experiments. We conducted training over 30 epochs across our comprehensive pairwise dataset, which includes 322 specimens. Additionally, separate models were trained on datasets 30A and 30B. The learning process was driven by an initial learning rate of 2×10^−4^ with the ADAM optimizer (Kingma and Ba, 2014), and was linearly reduced to 2×10^−6^ by the 30^th^ epoch with our learning rate scheduler.

As depicted in Figure 1, our training scheme begins with the estimation of new shape descriptors *F* and *G* for any pair of shapes within a dataset, specifically from the source and target shapes *S*_1_ and *S*_2_, respectively. Initially, these descriptors are derived from the wave kernel descriptors *D*_1_ and *D*_2_. Additional inputs for generating *F* and *G* include the precomputed Laplace and mass matrices (*L* and *M*), spatial gradient matrices (*G*), and the eigenvalues and eigenvectors of the Laplace-Beltrami operator (LBO) for both shapes. These computations are facilitated by our Siamese DiffusionNet function approximator. Further processing involves projecting *F* and *G* into the LB eigenbasis Φ and Ψ corresponding to shapes *S*_1_ and *S*_2_, respectively. This step is crucial for estimating the main branch functional maps via the regularized FMap block, as highlighted by the black lines in Figure 1. The projections also facilitate spatially and spectrally cycle consistent estimates from the cycle consistency block Π (depicted by the red in figure 1). Additionally, *F* and *G* are projected into the connection Laplacian basis for their respective shapes, which assists in estimating the complex functional correspondence *Q* using the Complex FMap Block depicted in orange in Figure 1.

**Figure 1-.**
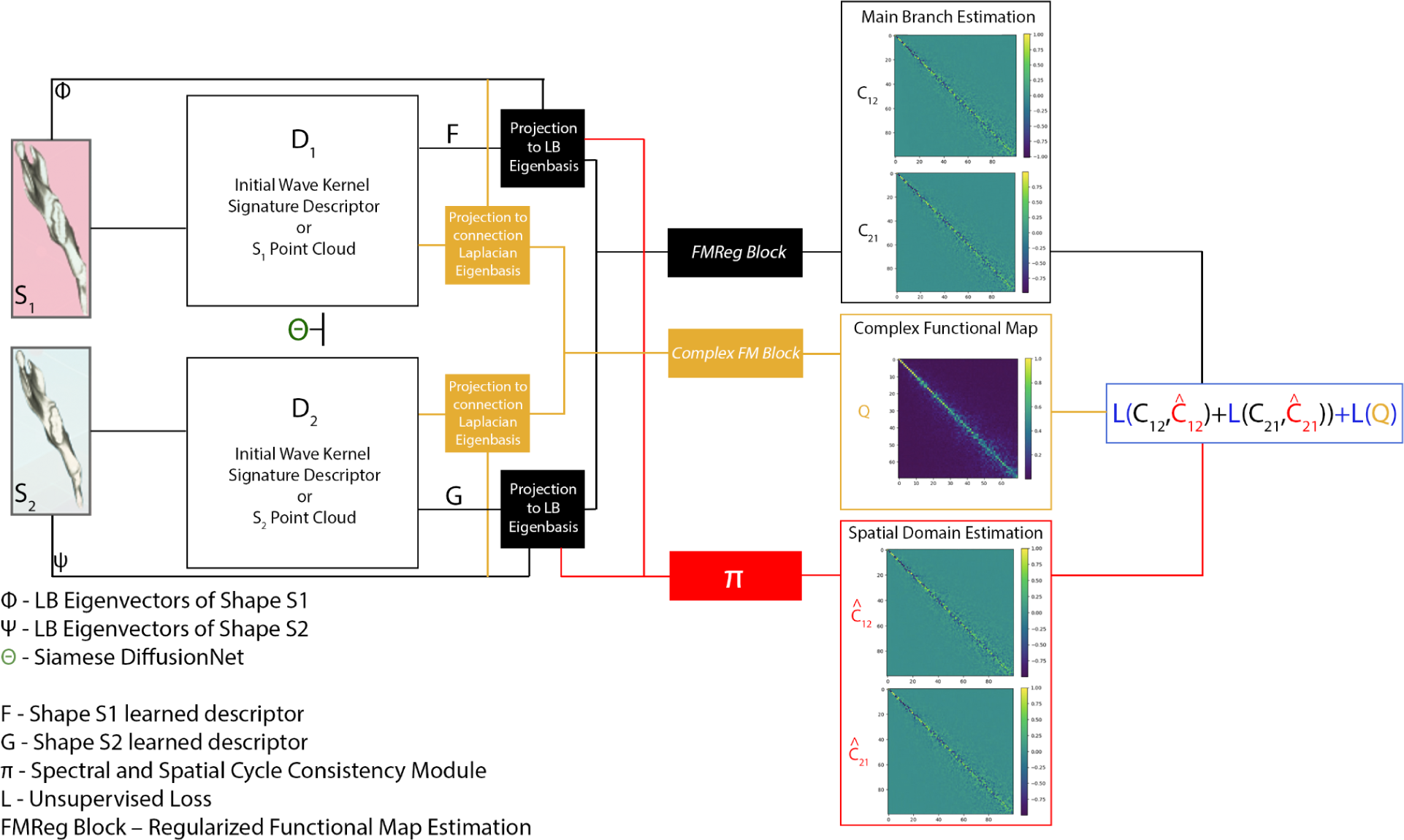
Network Architecture and Training Scheme. This figure illustrates the architecture of the enhanced MorphVQ pipeline developed for automatic landmark acquisition. The process begins with the acquisition of the Laplace-Beltrami (LB) eigenvectors (Φ, Ψ) for shapes S1 and S2, respectively. Each shape’s point cloud or wave kernel signature is processed through a Siamese DiffusionNet (Θ) to obtain improved shape descriptors (D1 for S1, D2 for S2). These descriptors (F and G) are then projected onto the Laplacian Eigenbasis for the main branch of the model, and the connection Laplacian eigenbasis for the complex functional map branch. The core of the pipeline, the FMReg Block, regularizes the functional map estimation, ensuring robust mapping, by integrating both spectral and spatial cycle consistency modules (Π), we obtain additional functional maps needed to compute the loss. The final output includes the complex functional map (Q) and estimation functional maps from the main branch and the cycle consistency branch in both directions (C_12 and C_21). The network optimizes these outputs by minimizing a combined loss function (L).

With Functional map estimates from both the FMReg Block 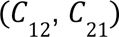 and the cycle consistency block Π 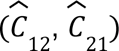 in both source-to-target and target-to-source directions and we are able to compute the first portion of our unsupervised loss as:

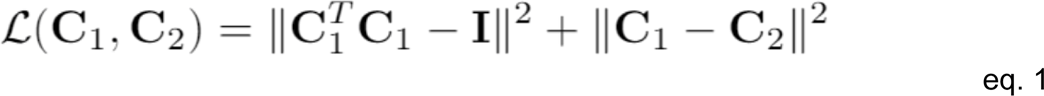

Here, the first term promotes the orthogonality of our main branch functional map, and the second term promotes the consistency of both maps from each branch. For additional information on how 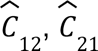 are obtained from the cycle consistency module Π see (Sun et al., 2023) section 4.1. The second portion of our unsupervised loss on the complex functional map *Q* is meant to impose orthogonal (Donati et al., 2022):

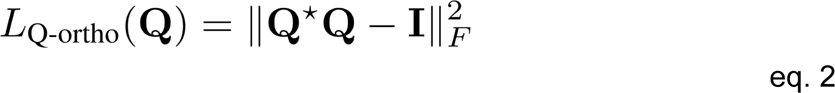

Jointly with the laplacian regularizer implemented in the Complex FM block (Figure 1) this loss in eq. 2 ensures that our model learns orientation-preserving isometric maps. With these losses computed given any pair of shapes, we compute the gradients with respect to the learnable parameters in our DiffusionNet function approximator and apply an update given a specific learning rate.

**Figure 2-.**
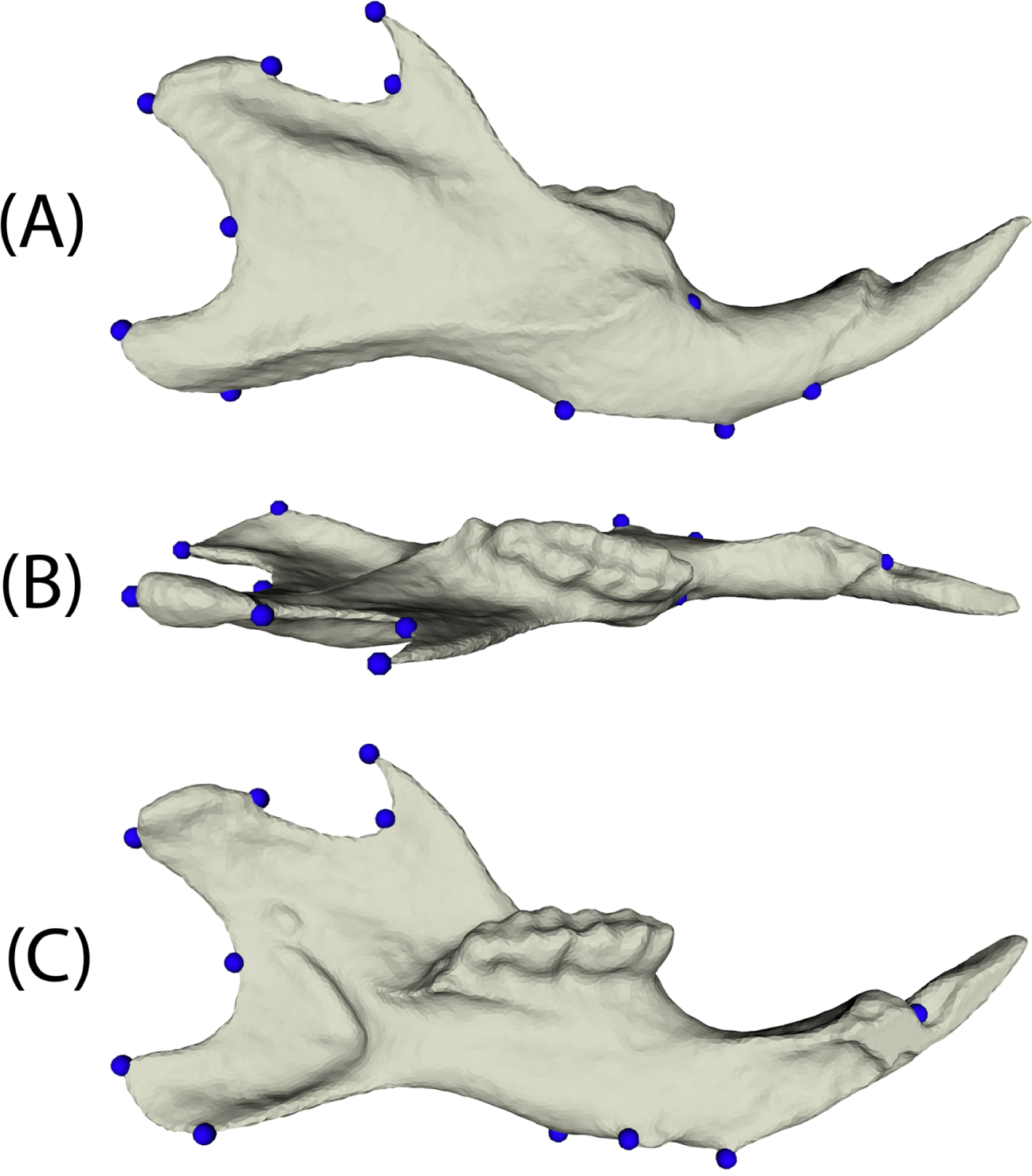
Mandibular landmarks used in the study. Three views of a landmarked mouse mandible. The landmarks are shown as blue spheres. (A) Lateral view in the sagittal plane presenting the side profile of the mandible, highlighting the overall shape and relative positions of the landmarks on the lateral surface. Key landmarks are located at the condylar process, the angular process, and the incisor region. (B) Dorsal view in the horizontal plane illustrating the superior aspect of the mandible, providing a clear view of the dental arcade and alignment of the teeth. (C) Mirrored medial view in the sagittal plane showing the inner side of the mandible.

### 2.4 Obtaining updated MorphVQ Landmark Estimates

After training the Deep Functional Mapping (DFMap) network within the updated MorphVQ pipeline, we conduct pairwise inference on all specimens used in unsupervised training, as well as on a separate dataset of 103 unseen (validation) specimens. We retain the functional maps (C_12_ and C_21_) and the Laplace-Beltrami Operator (LBO) eigenvectors for each pair of specimens. Dense point-to-point maps T_12_ are computed by projecting the eigenvectors of the source shape *S*_1_ onto the eigenspace of the target shape *S*_2_ using the functional map C_12_. The closest matches in this transformed space are identified using a 1-nearest neighbor search, thus deriving T_12_. Similarly, T_21_ is computed by reversing the roles of *S*_1_ and *S*_2_, thereby mapping each point on *S*_2_ to its nearest counterpart on *S*_1_.

To derive landmark estimates based on the dense point-to-point correspondences (T_12_ and T_21_) generated by our model, we utilized the VTK Python library (Schroeder et al., 2013). For each of the 13 ground-truth landmarks on every source shape, we identified the nearest vertex on the corresponding 3D model. The vertex index was then used to query T_12_ and T_21_, obtaining corresponding vertices on the respective target shapes. These target vertex coordinates were considered as the best estimates of the configuration of homologous landmarks on the target specimens. This procedure was repeated for each pair of source and target specimens within our pairwise dataset, providing *n*−1 estimates of landmark configurations for each ground-truth landmark set. From these *n*−1 estimates for each specimen, we selected the median set of landmark estimates as our primary estimated landmark configuration, as derived from our updated MorphVQ pipeline.

### 2.5 Estimating landmarks with MALPACA

As an alternative method of automatically landmarking the same set of samples, we used the MALPACA module in SlicerMorph (Zhang et al., 2022). MALPACA is the multi-template implementation of ALPACA, in which a 3D model of a landmarked specimen is used as a source model to transfer the same set of landmarks to target models using point-cloud based linear and deformable registrations (Porto et al., 2021). Multi-template version of ALPACA allows using more than one source model to generate multiple estimates of what the landmark position should be for the target sample. The median of these estimates were then used as the final estimate for the target, which reduces the potential bias from using a single reference and often provides estimates closer to the ground-truth (Zhang et al., 2022). For technical aspects of ALPACA pipeline, and its multi-template implementation we refer to the reader Porto et al., 2021 and Chi et al., 2022 respectively. For this study we used 7 randomly selected mandibles as reference models. We executed the MALPACA run on the same server as the geometric learning pipeline was run, but using only the CPU. During MALPACA, we used the Bayesian implementation of the Coherent Point Drift (https://github.com/ohirose/bcpd), which provides a significant speed up compared to the standard implementation (the acceleration option in the ALPACA menu). All other ALPACA settings were left as default. For clarity, MALPACA pipeline was not run on the mandibles derived from the preprocessing pipeline described in Section 2.1 above, but on the original mandibles models generated for Navarro and Maga 2016, as ALPACA does not require those preprocessing steps.

On average, final landmark positions for a target model were calculated from 7 templates in little less than 3.5 minutes; or in other words, obtaining a single ALPACA run for one template took less than 30 seconds. The full MALPACA pipeline for obtaining median estimates for all 425 samples was just over twenty four hours. MALPACA estimated landmarks and the ground-truth points they were compared to are provided in our github repository https://github.com/oothomas/SSC-MorphVQ.

### 2.6 Landmark error study

Once unscaled landmark estimates were obtained from both MALPACA and our updated MorphVQ, we employed the numpy library to calculate the root mean square error (RMSE) between our estimated points and the expert-measured ground-truth points. This calculation was averaged over three dimensions and performed for all 322 specimens in our main dataset, the 425 specimens in the MALPACA dataset, and for the landmark estimates derived from inference on the 103-specimen validation dataset for each of the three versions of the updated MorphVQ. These results are detailed in Table 2.

## 3. Results/Discussion

Here, we evaluate our model’s capacity to automatically and, in an unsupervised way, yield accurate landmarks, comparing its performance against MALPACA, the current state-of-the-art algorithm for automatic landmark placement. This critique includes a detailed error analysis, highlighting areas where our model outperforms the original MorphVQ.

### 3.1 Error-based Comparative Performance and Model Generalization

Our analysis focused on comparing the Root Mean Square Error (RMSE) of landmark predictions relative to manually placed ground-truth landmarks from our FMap-based models to those obtained from MALPACA, the state-of-the-art. The focus of our analysis is on the robustness of these FMap-based models, particularly in their ability to generalize to unseen specimens.

MALPACA generally exhibited the lowest Root Mean Square Error (RMSE) across all landmarks with an average RMSE of 0.113 mm, indicating high precision in landmark estimation on the full sample size. Despite this, our FMap-based models showed competitive performance, especially when considering that they were trained in an unsupervised manner on significantly smaller datasets (Table 1). Notably, the models trained on random selections of 30 specimens (30-Random A and B) performed comparably to the main model trained on 322 specimens in both training and validation phases. This suggests a strong generalizability of the FMap-based approach even with limited training data.

**Table 1-.** Root Mean Square Error Comparison across Datasets. This table presents a comprehensive summary of the Root Mean Square Error (RMSE) for landmark placement across various models and datasets. The RMSE values for each landmark (LM1 to LM13) and the overall mean RMSE are listed for the MALPACA model (using all samples), the main model (trained on 322 specimens), and two subset models (each trained on 30 random specimens from the main set). Results are divided into training and validation phases, allowing for comparison of model performance in learning and generalization contexts. The MALPACA model shows the lowest overall mean RMSE on all samples, while the subset models and main model exhibit competitive RMSE values, indicating robust performance even with reduced training sets.

| Model | Dataset | # Samples | RMSE Per-Landmark (mm) |  |  |  |  |  |  |  |  |  |  |  |  | RMSE |
| --- | --- | --- | --- | --- | --- | --- | --- | --- | --- | --- | --- | --- | --- | --- | --- | --- |
|  |  |  | LM1 | LM2 | LM3 | LM4 | LM5 | LM6 | LM7 | LM8 | LM9 | LM10 | LM11 | LM12 | LM13 | Mean |
| MALPACA - All Samples | - | 425 | 0.100 | 0.154 | 0.066 | 0.163 | 0.117 | 0.147 | 0.082 | 0.130 | 0.086 | 0.067 | 0.077 | 0.135 | 0.141 | 0.113 |
| Main Model | Training | 322 | 0.103 | 0.155 | 0.112 | 0.195 | 0.129 | 0.159 | 0.158 | 0.113 | 0.133 | 0.153 | 0.110 | 0.134 | 0.157 | 0.139 |
|  | Validation | 103 | 0.107 | 0.147 | 0.118 | 0.172 | 0.120 | 0.164 | 0.159 | 0.131 | 0.121 | 0.184 | 0.124 | 0.118 | 0.148 | 0.139 |
| 30-Random A | Training | 30 | 0.117 | 0.171 | 0.109 | 0.189 | 0.121 | 0.149 | 0.153 | 0.113 | 0.133 | 0.151 | 0.090 | 0.147 | 0.170 | 0.139 |
|  | Validation | 103 | 0.105 | 0.144 | 0.119 | 0.168 | 0.119 | 0.158 | 0.152 | 0.126 | 0.120 | 0.183 | 0.122 | 0.118 | 0.147 | 0.137 |
| 30 Random B | Training | 30 | 0.113 | 0.136 | 0.102 | 0.183 | 0.122 | 0.155 | 0.152 | 0.130 | 0.128 | 0.145 | 0.113 | 0.128 | 0.164 | 0.136 |
|  | Validation | 103 | 0.110 | 0.148 | 0.116 | 0.170 | 0.119 | 0.158 | 0.158 | 0.130 | 0.119 | 0.186 | 0.122 | 0.118 | 0.146 | 0.139 |

While MALPACA consistently showed low RMSE, the FMap-based models exhibited lower RMSEs for specific landmarks in certain instances. For example, landmarks LM3 and LM9 observed notably lower RMSEs in one or more of our models compared to MALPACA, highlighting instances where FMap models potentially outperform MALPACA in precision.

Interestingly, all FMap-based models, including those trained on just 30 specimens, maintained similar levels of performance when validated against the remaining 103 specimens not seen during training. This indicates not only the robustness of the FMap models in dealing with unseen data but also indicates their practical utility in scenarios where acquiring large volumes of training data is challenging or impractical.

**Figure 3-.**
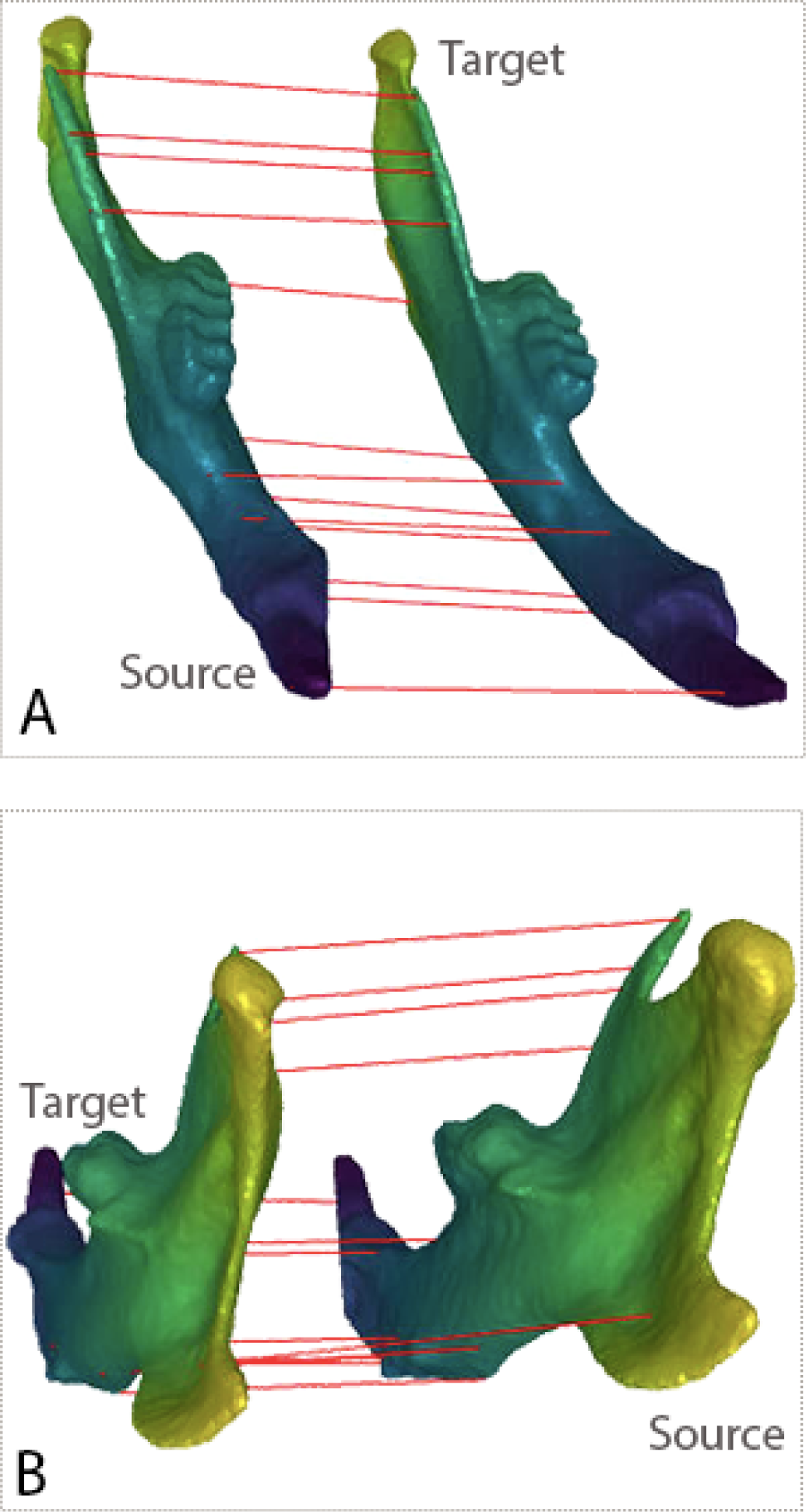
Point-to-Point Correspondence Quality. This figure shows the Point-to-Point correspondences obtained form our model for two randomly chosen mouse mandible specimen. Regions with the same color indicate point-to-point (morphological) correspondence

The distribution of RMSE values as visualized in the boxplots (Figure 4, 5, 6, and 7) corroborates these observations. Despite some variability and presence of outliers, the overlap in the interquartile ranges between our FMap models and MALPACA across several landmarks (e.g., LM4, LM7, LM10) suggests that our models are not only efficient in training but also effective in achieving high accuracy comparable to MALPACA. The ability of the FMap-based models to achieve similar accuracies to MALPACA has wide-ranging implications for the utility of this updated MorphVQ. Not only does it provide a more efficient alternative to existing methods but also extends the accessibility of high-precision landmark detection to studies with limited resources.

**Figure 4-.**
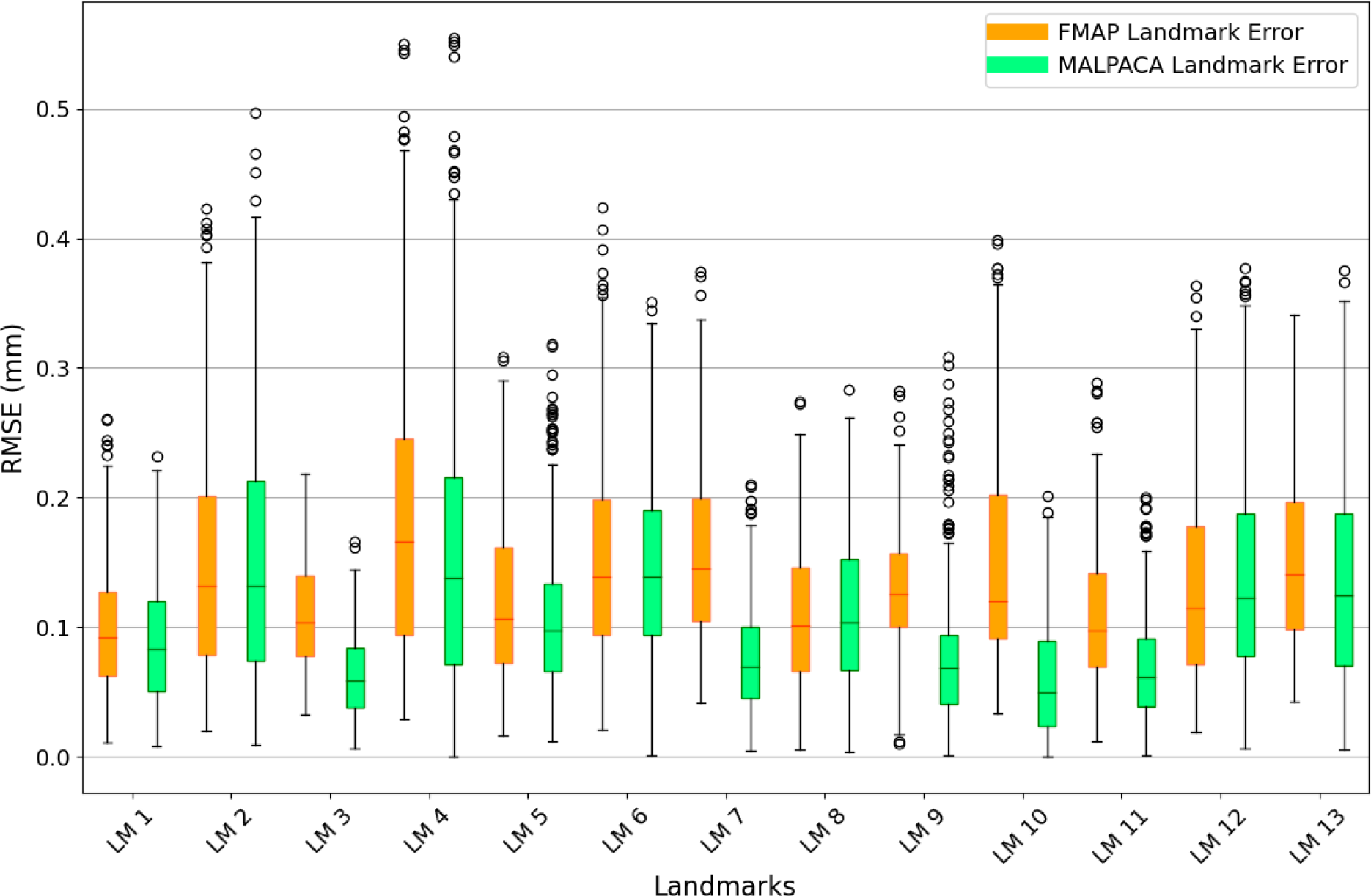
Comparison of Main Model to MALPACA Landmark Error Comparison. This figure presents a boxplot comparison of the Root Mean Square Error (RMSE) for landmark placement between our enhanced MorphVQ model (trained on 322 specimens) and the ground truth provided by the MALPACA algorithm. Each boxplot represents the distribution of RMSE values across different landmarks (LM1 to LM13) for both the FMAP and MALPACA methods. The orange bars indicate the RMSE for the FMAP method, and the green bars represent the RMSE for MALPACA, with outliers shown as circles. The plots illustrate the variance and median error for each landmark, providing a visual assessment of the precision and reliability of each method in landmark placement.

**Figure 5-.**
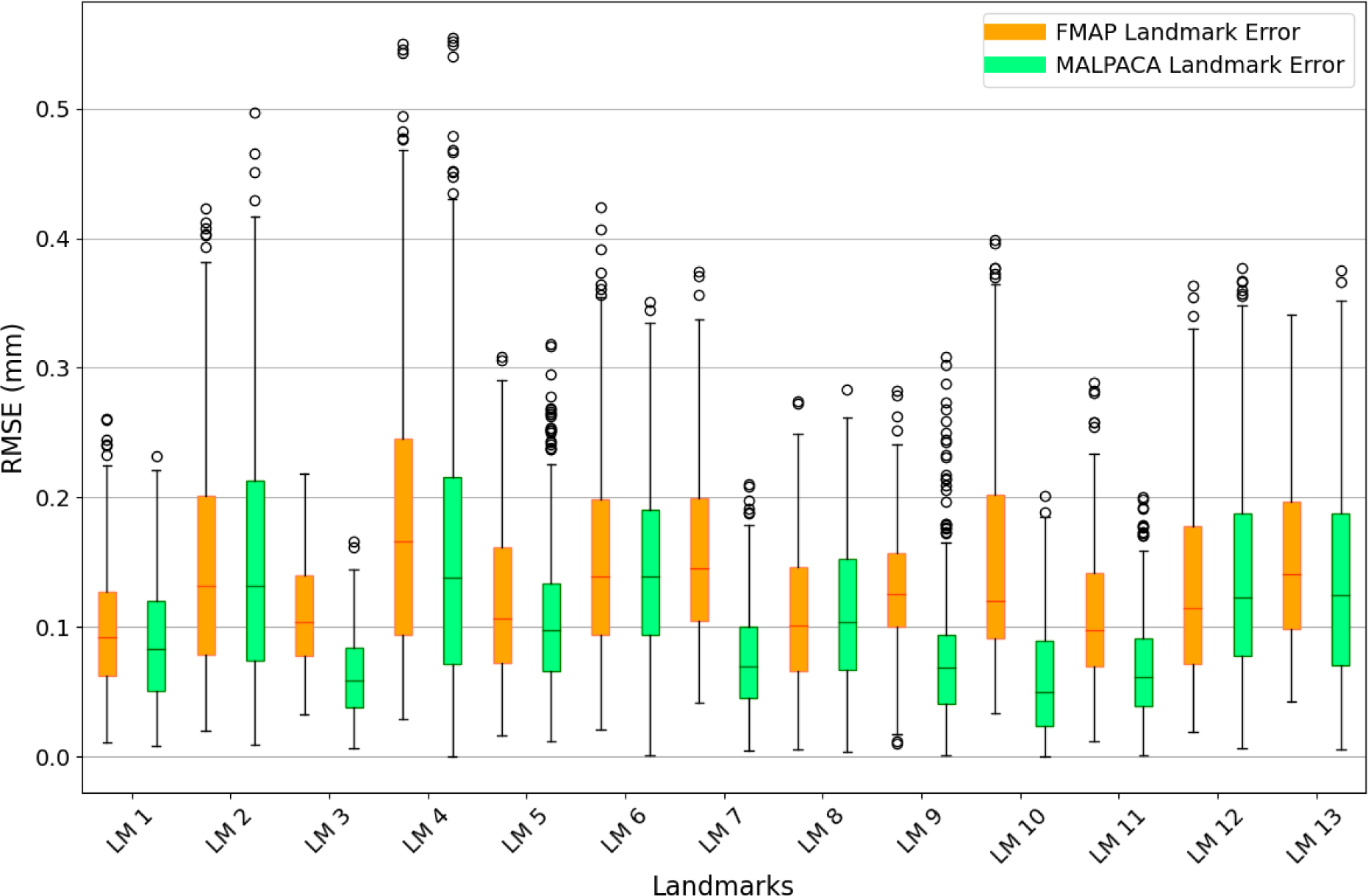
Main Model (Validation) to MALPACA Landmark Error Comparison. This figure displays a boxplot comparison of the Root Mean Square Error (RMSE) for landmark placement on 103 validation specimens, where the main model was used solely for inference. It compares the performance of our enhanced MorphVQ model (orange) against the established MALPACA algorithm (green) across thirteen landmarks (LM1 to LM13). This visual comparison illustrates the generalization ability of our model in accurately predicting landmarks in unseen data, highlighting its potential robustness and reliability in practical applications.

**Figure 6-.**
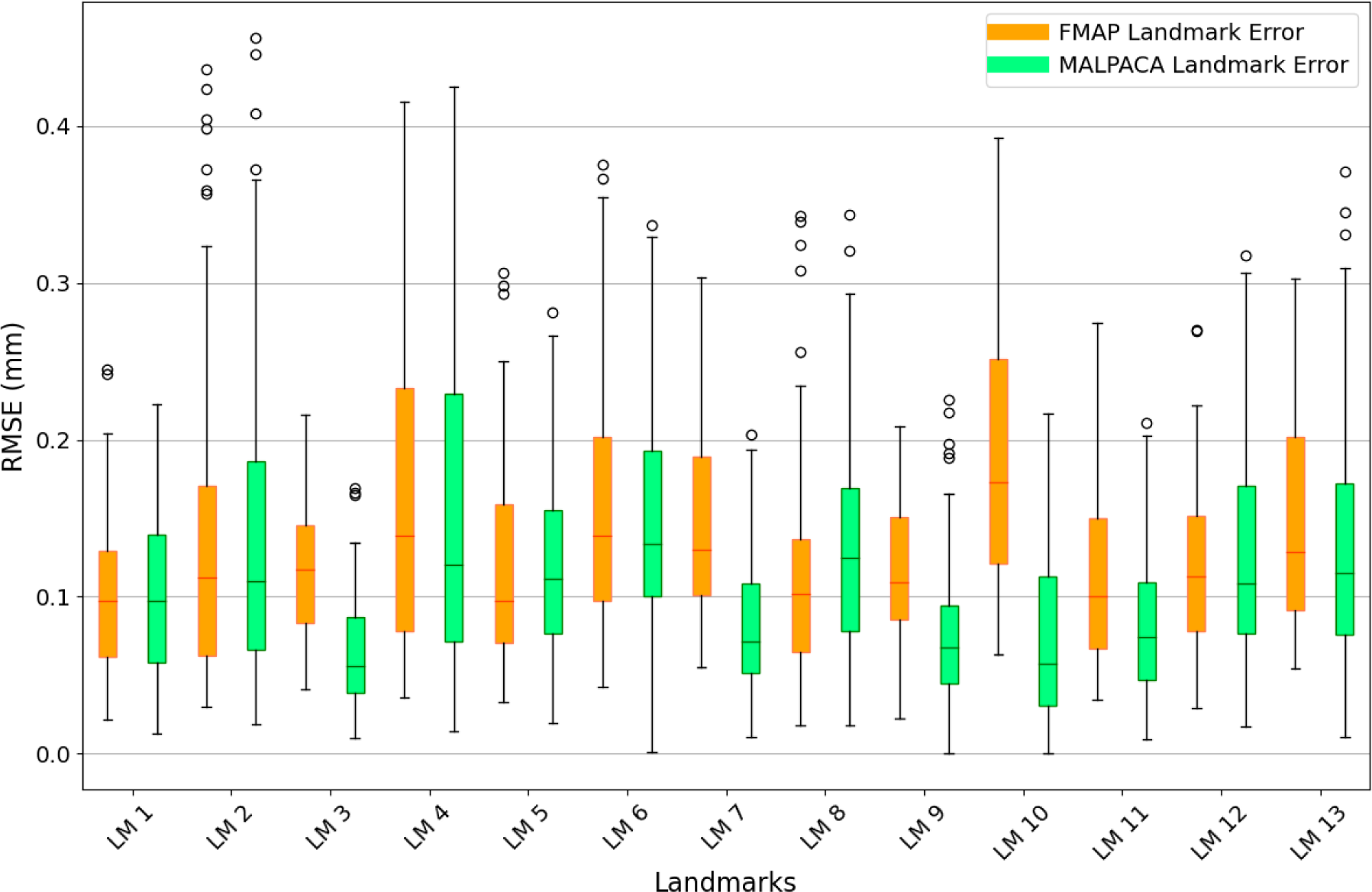
Model 30-Random A (Validation) to MALPACA Landmark Error Comparison. This figure depicts the Root Mean Square Error (RMSE) comparison for landmark placement using a second model, trained on 30 specimens selected from the original 322 used to train the main model. The comparison is made using the same 103 validation specimens. This visualization underscores the model’s effectiveness in generalizing from a smaller training set to unseen data, highlighting the potential scalability and efficiency of using reduced training sets in landmark placement applications.

**Figure 7-.**
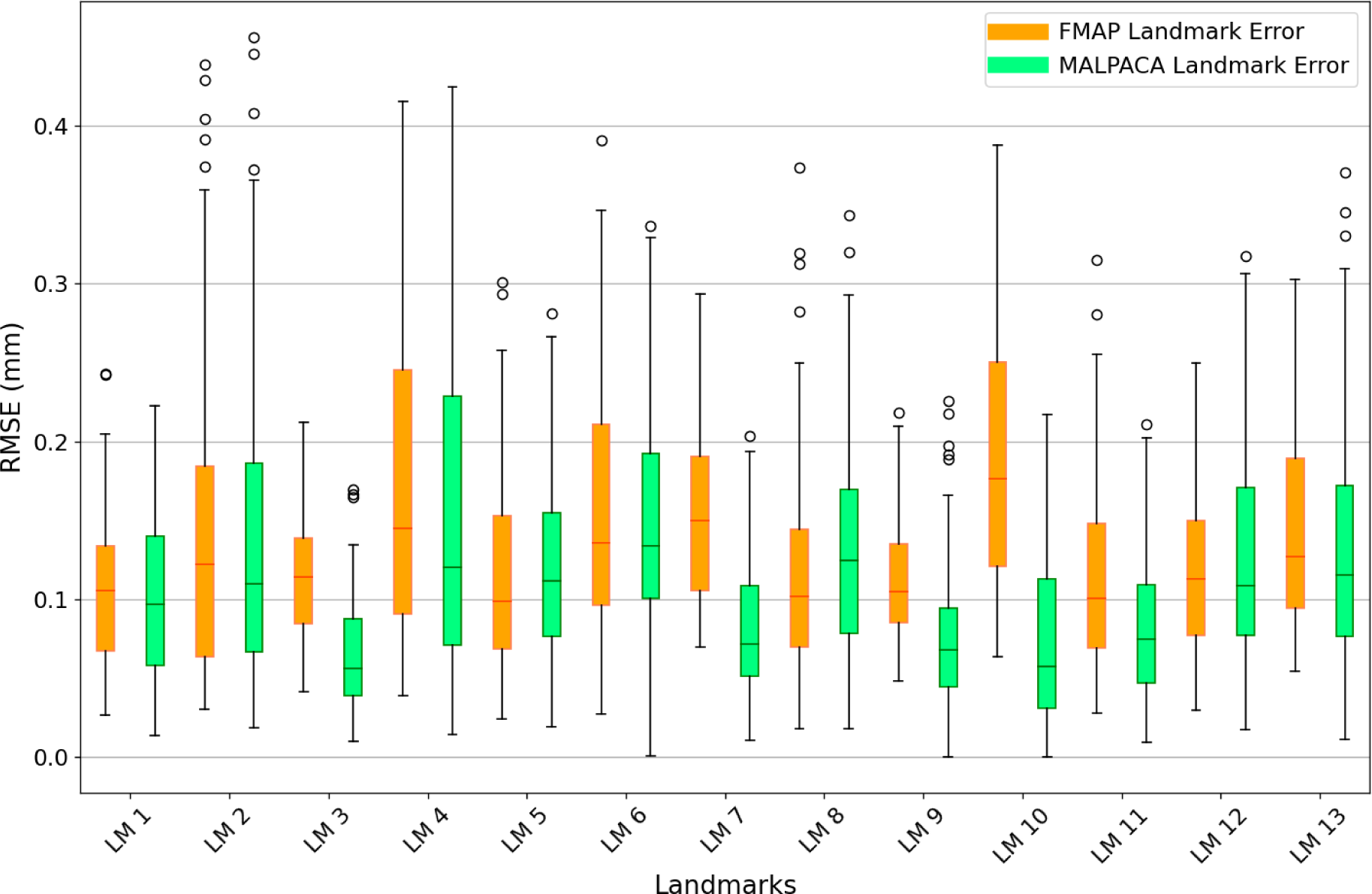
Model 30-Random B (Validation) to MALPACA Landmark Error Comparison. This figure shows the Root Mean Square Error (RMSE) comparison for landmark placement based on a third model, trained on a different subset of 30 specimens from the original 322 used to train the first and validated solely on the same 103 specimens used previously. This analysis highlights the model’s robustness and consistency in generalizing from different small training sets to the same validation set, underscoring the potential for using smaller, diverse training subsets to achieve reliable landmark placement.

The findings underscore the potential of unsupervised learning models in anatomical landmark detection and placement, suggesting that they can offer a viable alternative to more traditional, supervised methods in certain applications. The scalability and efficiency of these models make them particularly suited for large-scale morphometric studies where rapid processing and generalization to new specimens are crucial.

### 3.2 Efficiency and Correspondence Quality of the new MorphVQ

Our updated MorphVQ demonstrates significant efficiency advantages over MALPACA in both training and inference phases. We found that training the model on the entire dataset of 322 specimens was unnecessary, as models trained on just 30 specimens performed equivalently. Additionally, these 30-specimen models achieved convergence within five epochs, rendering the full 30 epochs redundant. This reduction in training duration does not compromise the model’s effectiveness. Specifically, the model trains on a dataset of 30 specimens, each with 12,000 vertices processed pairwise, in approximately three hours. Inference on the remaining 395 specimens, also processed pairwise, takes about 12 hours. This is significantly faster than MALPACA, which would have taken 22 hours to generate landmarks for the same 395 specimens.

For a fairer comparison between our updated MorphVQ and MALPACA, the preprocessing steps required by each approach must be considered. In both cases, mandibles must be segmented properly to isolate them from the rest of the tissue in the microCT scan. While both approaches require generating polygon models for each specimen, the updated MorphVQ approach demands manifold polygon models and additional preprocessing that is not required for ALPACA. A manifold polygon mesh is a type of 3D model where the geometry is well-defined such that each edge belongs to exactly two faces, ensuring that the mesh represents a continuous surface without holes or self-intersections. This property is crucial for many computational algorithms, including those used in functional map-based analysis. This additional preprocessing can be automated in 3D Slicer with the help of Python libraries such as pyMeshFix, as described in our methods (see the pre-processing directory of our repository at https://github.com/oothomas/SSC-MorphVQ for additional details). This process took approximately 1.5 hours to produce meshes with the properties needed for FMap-based analysis.

Additionally, both approaches require landmarking a small subset of specimens to use as templates for generating landmarks, but for very different purposes. In MALPACA, these landmarks are collected before the analysis. In our approach, these landmarks can be chosen after training and are used to query the pairwise point-to-point maps for corresponding landmarks across all specimens in the dataset, and as such, if landmark configurations need to be modified (e.g., by adding new landmarks), it can be easily accomplished by re-querying the point-to-point map for the new configuration. Our approach yields full point-to-point correspondences between all pairs of shapes. As a result, there is no need to re-train the model to obtain various landmark configurations representing different biological hypotheses; the full point-to-point correspondences are always valid and reusable. In contrast, in the current implementation of ALPACA, a new landmark configuration requires rerunning the algorithm with new landmarks. Although, the current implementation of ALPACA can be revised to achieve the same functionality.

The efficacy of our method is highlighted by the high quality of the FMaps and point-to-point correspondences it produces. For example, Figure 3 illustrates the bijectivity between a randomly selected pair of shapes, with similar colors on the source and target shapes indicating corresponding anatomical or morphological regions. This high level of correspondence has been consistently observed across all shape pairs that we have manually reviewed. Although a complete manual review of all shape pairs is impractical due to the sheer number, the sampled checks confirm the model’s reliability and accuracy. Notably, these point-to-point correspondences were obtained without any post-processing refinement, which was essential in the original MorphVQ. By incorporating spatial and spectral cycle consistency during training, this step is now unnecessary, enhancing correspondence quality and reducing the overall processing time of the pipeline.

### 3.3 Visual Evaluation of Landmark Placement Accuracy

The precision of landmark placement on mouse mandibles by both the MALPACA and our Functional Map (FMap)-based models was investigated with their suitability for quantitative trait loci (QTL) analyses in mind. Despite MALPACA exhibiting a marginally lower Root Mean Square Error (RMSE) on average across all landmarks, both methodologies displayed a high level of precision that meets the requirements for the intended applications.

As depicted in Figure 8, the landmarks placed by our FMap-based approach (green spheres) generally coincide with those identified by the ground-truth (blue spheres) and MALPACA (red spheres). This visual congruence reinforces the utility of our model, particularly in the context of the biological studies for which these landmarks are critical. Notably, while there are deviations in a few cases, these remain within an acceptable range for our purposes, which is impressive considering that high precision is critical for our particular morphometric application.

**Figure 8-.**
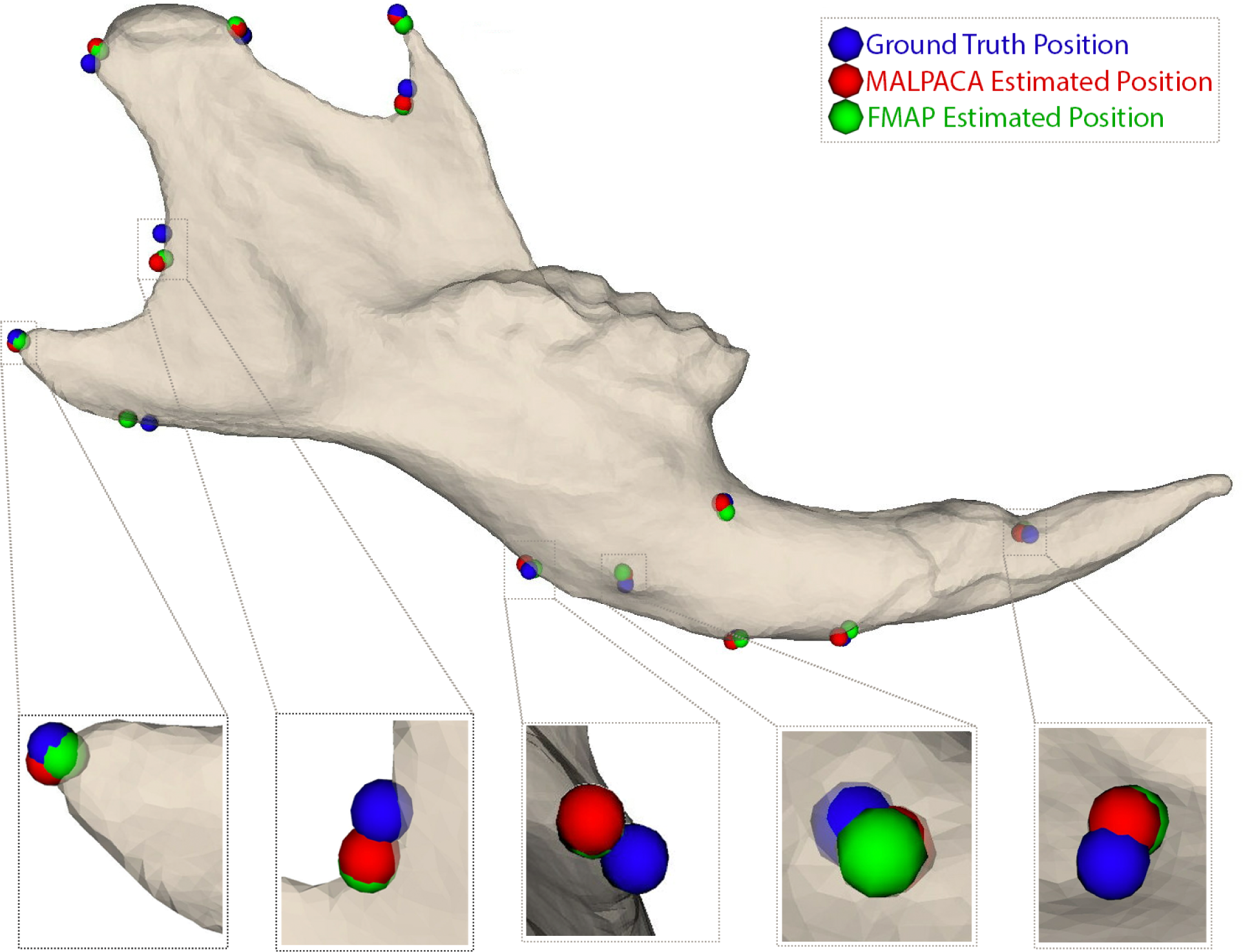
Visual Comparison of Landmark Placement Accuracy on a Mouse Mandible. This figure presents a visual comparison of landmark placement on a mouse mandible, illustrating the ground truth positions (blue), MALPACA estimated positions (red), and estimates from our FMAP model (green). Insets zoom into selected landmarks to highlight the proximity of the estimated positions to the ground truth. Despite the slightly lower Root Mean Square Error (RMSE) associated with MALPACA, the landmarks estimated by our FMAP model align closely with both the ground truth and MALPACA’s estimates, demonstrating high-quality landmark estimation. This visual representation underscores that the deviations between the methods are minimal and generally fall within the range expected from human error in manual landmark placement. Thus, the FMAP model offers comparable precision, reinforcing its utility and reliability in morphometric analyses.

One of the most significant features of the updated model is the generation of full point-to-point correspondence maps for all pairs of specimens. This functionality enables researchers to select any landmark configuration from the dataset, facilitating flexibility in testing various hypotheses or adapting the model to different applications. Such versatility is particularly beneficial in dynamic research environments where analytical needs can evolve over the course of a study.

Also, our FMap-based approach generates comprehensive point-to-point correspondences by considering correspondences between all pairs of shapes in the dataset. This method offers a robust alternative to template-based methods, which may introduce bias by relying on landmark estimations from a limited number of specific template specimens. In future studies, the choice between these methods should account for the precision requirements, potential bias in landmark estimation, and the available sample size for landmarking.

### 3.4 A Refined MorphVQ Model

The model we introduce is a refined version of the MorphVQ model, incorporating significant enhancements over its predecessor and addressing key limitations in the shape matching portion of the MorphVQ pipeline.

The first improvement is its orientation-aware capability, which eliminates the need for approximate rigid pre-alignment of specimens. Originally, the MorphVQ pipeline required rigid pre-alignment using the Auto3DGM algorithm, which was time-consuming for large datasets. Our study integrates the Complex FMap method, which uses complex-valued functions to align tangent vector fields, bypassing the need for pre-alignment and streamlining the workflow. This approach reduces issues such as local and global symmetry flipping, preserves global orientation, and enables an unsupervised learning framework without the need for supervised loss functions or reliance on extrinsic descriptors (Donati et al., 2022).

The second improvement is the adoption of the DiffusionNet model (Sharp et al., 2020) for descriptor learning, replacing the Harmonic Surface Network (Wiersma et al., 2020). DiffusionNet is designed for deep learning on 3D surfaces, using diffusion processes for effective spatial information transfer. It overcomes limitations of traditional convolutional networks on non-flat geometries, achieving robustness against variations in resolution and surface sampling. DiffusionNet’s ability to learn across different geometric representations allows training on point clouds from polygon mesh models, making it suited for establishing non-rigid correspondences among variable biological 3D shapes. Its integration of learned diffusion processes with spatial gradient features simplifies the learning process and enhances performance in unsupervised learning settings for deformable surfaces.

The third and most important innovation in MorphVQ is based on the work of Sun et al. (2023) to enforce spatial and spectral cycle consistency during the Deep Functional Maps (DFMaps) network training process. Cycle consistency is essential for non-rigid shape matching as it ensures that mappings between shapes remain consistent both in feature-based and point-wise domains when cycled through a series of shapes. This robust regularizer optimizes mappings across a collection of shapes.

Traditionally, DFMaps project shape features into a spectral domain, using eigen decompositions related to the shape’s geometry. While these spectral mappings can be consistent when cycling through shapes, they do not necessarily guarantee point-wise consistency, resulting in lower quality point-to-point correspondences and ultimately flawed landmarks. In the original MorphVQ, cycle consistency was used to improve mappings in a post-processing step (Huang et al., 2020). However, the new approach incorporates this during the training process, enhancing accuracy and reducing the need for additional refinement.

These advancements in the revised MorphVQ model represent a significant leap forward for automated anatomical landmark placement. By reducing the need for extensive preprocessing, enhancing model generalization, and increasing computational efficiency, this model sets a new standard for automatic landmark detection. It integrates modern computational methods with established morphometric practices, reaffirming the relevance of Geometry Processing and FMap-based methods in biological shape analysis while opening new opportunities for detailed anatomical studies.

## 4. Conclusions

In this study, we addressed the challenges of precise landmark acquisition on morphological datasets through the development and evaluation of an enhanced MorphVQ pipeline. Building on foundational techniques in geometry processing, our approach has successfully leveraged Functional Map correspondences to enhance the accuracy and efficiency of automatic landmark placement on morphological specimens represented as polygon meshes.

Our findings indicate a marked improvement in the precision and reproducibility of landmark placement. The integration of the DiffusionNet model, which harnesses the robustness of diffusion processes across complex geometries, along with the implementation of orientation-preserving complex functional maps, has substantially reduced the need for pre-alignment. This not only streamlines the workflow but also preserves the integrity of morphological data by avoiding distortions typically introduced by errors during rigid pre-alignment steps. The efficacy of these enhancements was empirically validated, demonstrating that our approach could achieve landmark placement accuracy with a Root Mean Square Error (RMSE) competitive with, and in some instances superior to, the current state-of-the-art method, MALPACA.

Moreover, the incorporation of spatial and spectral cycle consistency has significantly improved the bijectivity of mappings across highly variable biological shapes. This improvement is evidenced by the fact that no post-processing refinement of our correspondences between shapes was necessary, as our new MorphVQ provides high-quality functional and point-to-point maps without that final step.

This study not only demonstrates the viability of our updated pipeline in achieving high precision in landmark placement but also sets the stage for future research that might explore new computational techniques or refine existing ones to further enhance the capabilities of morphometric analysis. As we continue to unravel the complex narratives of biological evolution, the tools we refine today will undoubtedly play a pivotal role in the morphological studies of tomorrow.

## 5. Acknowledgments

This research was funded by grants from the National Science Foundation (OAC-2118240, HDR Imageomics Institute) and National Institutes of Health (NICHD-HD104435) to AMM. SlicerMorph tools and the data used in this research were previously funded by grants to AMM from the National Science Foundation (DBI 1759883) and the National Institute of Craniofacial and Dental Research (DE021417). I would like to thank my colleagues at the Seattle Children’s Hospital Research Institute, Dr. Rachel Roston and Dr. Sara Rolfe, for their feedback on my manuscript. I also extend my gratitude to my graduate school advisors, Dr. Mark Hasegawa-Johnson and Dr. John D. Polk, for their valuable input on the manuscript. Additionally, I appreciate the guidance and support of my undergraduate advisors, Dr. Ryan L. Raaum and William H.E. Harcourt-Smith. I am particularly grateful to Dr. Maks Ovsjanikov of the Ecole Polytechnique, France, for his initial advice and guidance regarding the methodological approach employed in this work.

## 6. Code and Data availability

The source code and the data to reproduce the findings of this study can be found in https://github.com/oothomas/SSC-MorphVQ, along with instructions on how to prepare the computing environment necessary for training and initiate the model.

